# ELI trifocal microscope: A precise cryogenic fabrication system to prepare target cryo-lamellae of cells for *in situ* cryo-ET study

**DOI:** 10.1101/2022.08.15.503951

**Authors:** Shuoguo Li, Ziyan Wang, Xing Jia, Tongxin Niu, Jianguo Zhang, Guoliang Yin, Xiaoyun Zhang, Yun Zhu, Gang Ji, Fei Sun

## Abstract

Cryo-electron tomography (cryo-ET) has become a powerful approach to study the high-resolution structure of cellular macromolecular machines *in situ*. However, the current correlative cryo-fluorescence and electron microscopy lacks sufficient accuracy and efficiency to precisely prepare cryo-lamellae of target locations for subsequent *in situ* cryo-ET structural study. Here, we developed a precise cryogenic fabrication system, the ELI trifocal microscope (ELI-TriScope), by setting up an electron (E) beam, a light (L) beam and an ion (I) beam at the same focal point to achieve accurate and efficient preparation of a target cryo-lamella without sacrificing the throughput. ELI-TriScope was developed starting from a commercial dual-beam scanning electron microscope (SEM) by incorporating a cryo-holder-based transfer system and embedding an optical imaging system just underneath the vitrified specimen. Cryo-focused ion beam (FIB) milling can be accurately navigated by monitoring the real-time fluorescence signal of the target molecule. Using ELI-TriScope, we prepared a batch of cryo-lamellae of HeLa cells targeting the centrosome, with an ∼100% success rate, and discovered new *in situ* structural features of the human centrosome through a subsequent cryo-ET structural study.

## Introduction

“ Seeing is believing” ; three-dimensional (3D) visualization of the cellular ultrastructure is an important step in understanding life. With the rapid technology development, cryo-electron microscopy (cryo-EM) single-particle analysis has become one of the most important tools to study 3D high-resolution structures of biomacromolecules *in vitro*^1-3^. Meanwhile, cryo-electron tomography (cryo-ET) has also been rapidly developed, becoming a unique technique to study the *in situ* high-resolution structures of biomacromolecular complexes and the locations and interactions of these complexes with their native cellular environment^4^, which will be expected to bring another revolutionary breakthrough in structural biology in the near future^5-8^.

However, many obstacles still exist in applying cryo-ET widely and efficiently for *in situ* structural study, especially the low quality, low efficiency, and low accuracy of specimen preparation methods^9-15^. Due to the limited penetration distance of electrons at the current accelerating voltage (300 kV) of the modern microscope, a thin (∼ 200 nm) cryo-section of a cell or tissue specimen should be prepared. Cryo-focused ion beam (cryo-FIB) milling has been proven to be an efficient method to prepare high-quality cryo-lamellae of cells for *in situ* structural study^16^, with many successful applications^17-22^. The lamella can be milled to 50∼300 nm thickness without the conventional artefacts caused by cryo-ultramicrotomy^23,24^. Recently, 3D architectures of various cellular organelles, such as the cytoskeleton^17^, endoplasmic reticulum (ER)^25^, and 26S proteasome^26^, have been revealed *in situ* by cryo-FIB and cryo-ET.

Furthermore, to enable accurate preparation of cryo-lamellae in the target region by cryo-FIB milling, the cryo-correlative light and electron microscopy (cryo-CLEM) technique has been developed. The target region in the cell can be labelled with a fluorescence molecular probe (e.g., GFP) and imaged by various cryo-fluorescence microscopies (cryo-FMs). Then, the fluorescence image can be used to navigate cryo-FIB fabrication. The conventional cryo-CLEM workflow requires a stand-alone fluorescence microscope with a cryo-stage for cryo-fluorescence imaging^11,12,14,27,28^, followed by cryo-scanning electron microscopy (cryo-SEM). A correlation alignment between the cryo-FM and cryo-SEM images is generated and used to guide cryo-FIB milling^15,29-31^. This workflow is complicated, and ensuring the sample quality is challenging due to multiple transfers between microscopes^30,32^. Each transfer increases the risk of sample devitrification and ice contamination, leading to reduced accuracy of the correlation alignment^33-35^. Moreover, specific fiducial markers imaged by both cryo-FM and cryo-FIB, such as fluorescent beads, are required for the correlation alignment using specific 3D correlative software^30,36^. Accurate z-axis positions of both fiducial markers and fluorescent targets are necessary for precise correlation; however, this information can only be roughly determined by widefield fluorescence microscopy^30,35^ or cryo-Airyscan confocal microscopy (CACM)^29^ with essentially limited resolution^15,37,38^.

Recently, another cryo-CLEM concept has been developed by integrating a fluorescence imaging system into the cryo-FIB/cryo-SEM chamber to avoid specimen transfer during microscopy^39^, and the commercially available products include iFLM^40^ (ThermoFisher Scientific, USA) and METEOR^41^ (Delmic, Netherlands). Using these integrated systems, cryo-FM images can be acquired before and after cryo-FIB milling to check the presence of the target signal without increasing the risk of contamination. However, these current systems potentially have a correlation resolution limitation due to the low numerical aperture (NA) of the objective lens, and the cryo-FIB milling efficiency is limited due to the frequent switching between cryo-FM and cryo-FIB.

Here, starting from the concept of our previously developed high-vacuum optical platform for cryo-CLEM (HOPE)^28^, we developed a new cryo-CLEM system named ELI-TriScope to achieve accurate and efficient preparation of target cryo-lamellae. Based on a commercial dual-beam scanning electron microscope (SEM), ELI-TriScope incorporates a cryo-holder-based transfer system and embeds an inverted fluorescence imaging system just underneath the vitrified specimen. In ELI-TriScope, an electron (E) beam, a light (L) beam and an ion (I) beam are precisely adjusted to the same focal point; as a result, cryo-FIB milling can be accurately navigated by monitoring the real-time fluorescence signal of the target molecule. With ELI-TriScope, there is no need to add fiducial markers or perform sophisticated correlation alignment between cryo-FM and cryo-SEM, leading to essentially improved efficiency, accuracy, success rate and throughput of cryo-FIB milling compared with other reported cryo-CLEM techniques.

To evaluate the efficiency of ELI-TriScope, the human centriole was selected as a challenging target to perform an *in situ* structural study. A batch of cryo-lamellae of HeLa cells targeting the centrosome were efficiently prepared with an ∼100% success rate. The subsequent cryo-ET study not only confirmed the ultrastructure of various typical components in human centrioles but also revealed new structural features. Therefore, ELI-TriScope provides a highly successful solution for sample preparation for *in situ* cryo-ET study and will have wide application in future *in situ* structural biology.

## Results

### Design of the ELI-TriScope system

Our ELI-TriScope system is developed based on a dual-beam SEM and contains two major components, a custom-designed cryo-holder-based vacuum transfer system for SEM imaging and an inserted fluorescence imaging system just underneath the vitrified specimen inside the SEM chamber (Fig. 1a and Extended Data Fig. 1; Supplementary Video 1).

**Figure 1.**
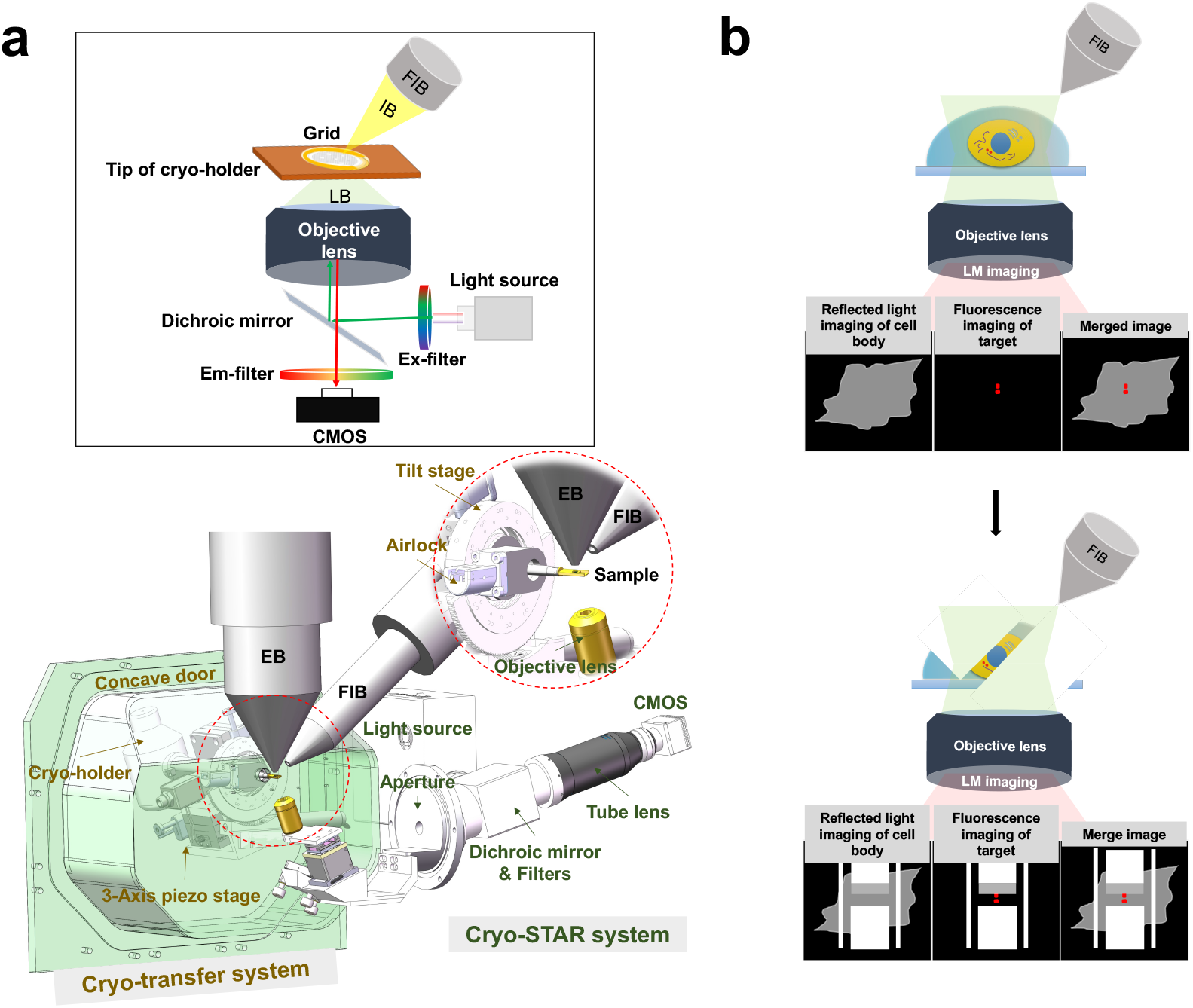
Design and principle of ELI-TriScope. (**a**) Schematic diagram and design drawing of ELI-TriScope in its operational mode. Each part of the system is labelled and described. IB, ion beam. LB, light beam. EB, electron beam. FIB, focused ion beam. (**b**) Working principle of ELI-TriScope. The ion beam and light beam are simultaneously focused on the area of interest of the cryo-specimen. The reflected light image recorded by the camera displays the cell body, and the target signal in the fluorescence image is used to navigate cryo-FIB milling.

For the vacuum transfer system, the original chamber door of the SEM is replaced by a new concave door equipped with an outside 3-axis piezo (New Focus 9161M, Newport, California) and servomotor (Maxon, Switzerland) stage tightly mounted with a cryo-holder adaptor (Extended Data Fig. 1a-b). The 3-axis stage allows high-precision movement in three dimensions. A worm wheel mechanism is designed to drive the 3-axis stage to tilt in the range of -70 to 55 degrees by the servomotor (Maxon, Switzerland) with a precision of 0.01 degrees. A homemade airlock prepump vacuum system developed previously^28^ connects the cryo-holder adaptor and the concave door (Fig. 1a). The cryo-holder can be loaded into the SEM chamber via the adaptor and through the prepump vacuum system. Then, the position of the cryo-holder can be adjusted by controlling the 3-axis stage, which allows precise localization of the cryo-specimen at the crossover point of the electron and ion beams. The movement and tilt of the stage can be controlled via customized software written in LabVIEW (National Instruments, USA).

Inside the SEM chamber, a widefield optical imaging system is inserted just underneath the tip of the cryo-holder (Fig. 1a and Extended Data Fig. 1a-c; see also Methods). We named this optical imaging system the <u>cryo</u>genic <u>s</u>imul<u>ta</u>neous monito<u>r</u> (cryo-STAR) system. To maximize the detection sensitivity for weak fluorescence signals of target molecules in real time, a high-NA dry objective lens is selected and installed in this system. Meanwhile, an epifluorescence system with a full-spectrum (DAPI/GFP/RFP/Cy5) white LED light source is equipped. The fluorescence signals from the vitrified specimen are collected by the objective lens and recorded by a high-sensitivity complementary metal–oxide–semiconductor (CMOS) camera.

The ELI-TriScope system adjusts the electron beam, ion beam and light beam to the same focal point. Therefore, the fluorescence signal of target molecules can be monitored in real time while cryo-FIB milling is being performed (Supplementary Video 1). As a result, the cryo-FIB fabrication procedure can be accurately navigated to the specific region of interest (Fig. 1b). Since there is no need for specimen transfer, the risk of ice contamination, specimen damage and devitrification can be largely avoided, and the operation time of fabricating one cryo-lamella can be efficiently reduced.

### ELI-TriScope workflow

The centrosome is a highly ordered organelle that controls cell proliferation, motility, signalling and architecture^42,43^. As the microtubule (MT)-organizing centre (MTOC) of vertebrate cells, centrosomes organize the formation of two poles of the mitotic spindle during cell division and act as templates for the formation of flagella and cilia^44-46^. Each centrosome in a cell consists of an MT-based cylindrical component named the centriole and a surrounding pericentriolar material (PCM)^46^. Most of the time, there are only two centrioles closely localized in one cell, and these two centrioles are termed the mother and daughter according to their maturity degrees. Therefore, preparing a cryo-lamella containing centrioles for *in situ* structural study is very difficult using a conventional cryo-FIB procedure without precise fluorescence signal navigation. We therefore select the human centriole as a challenging target to validate the performance of our ELI-TriScope technique with the following workflow (Fig. 2 and Supplementary Video 2).

**Figure 2.**
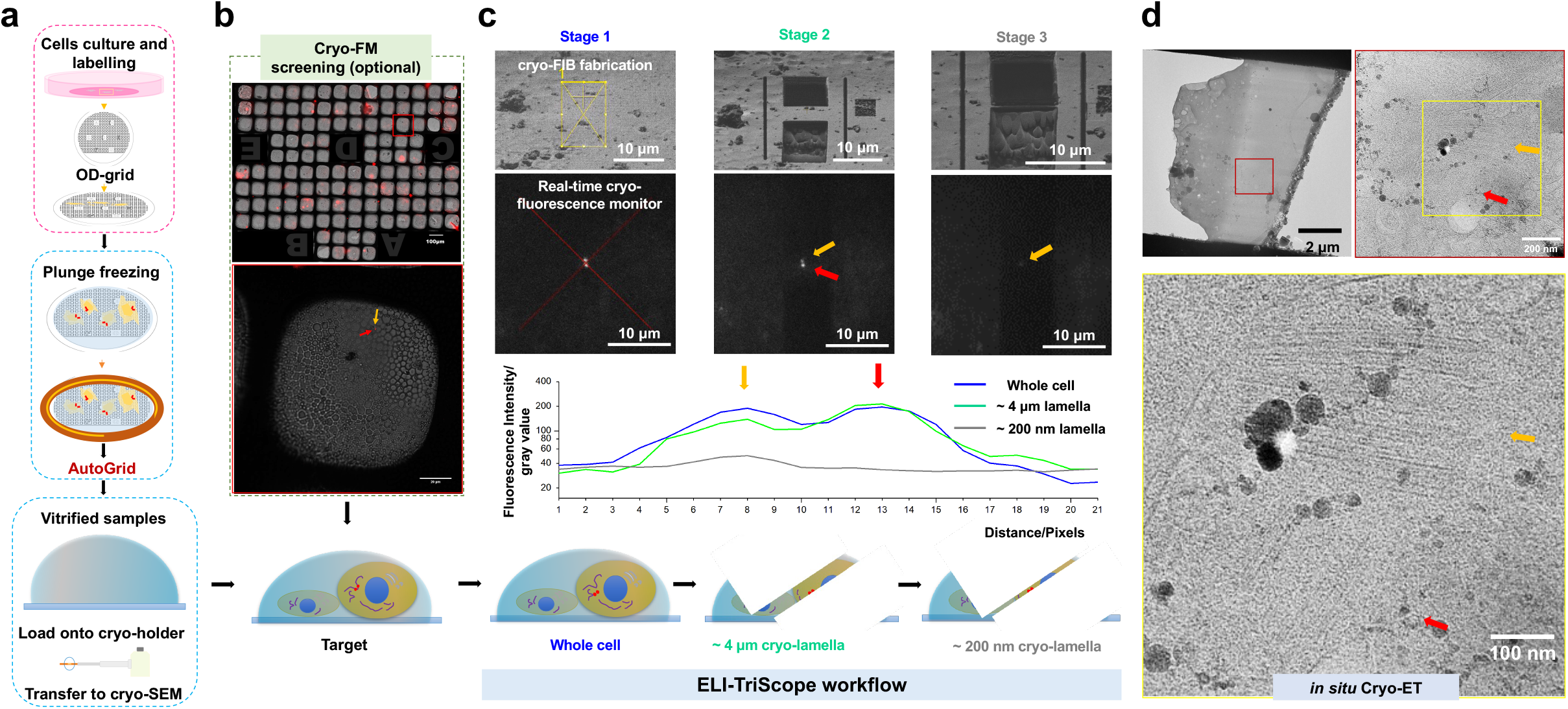
Cryo-CLEM workflow using ELI-TriScope. (**a**) Cells labelled by fluorophores are cultured on an OD grid (T11012SS, TIANLD, China), loaded into an FEI AutoGrid (ThermoFisher Scientific, USA), vitrified by plunge freezing and then loaded onto a cryo-holder. (**b**) Screening of cryo-vitrified cells using cryo-FM (optional). The ice thickness and the fluorescence signal are checked at this step. The bright-field atlas merged with fluorescence images of the grid can be acquired, and the positions of cells with good fluorescence signals (orange and red arrows) are recorded. (**c**) Real-time fluorescence signal-navigated cryo-FIB milling by ELI-TriScope. During the cryo-FIB process, the cryo-fluorescence intensity of the target is monitored in real time. In the SEM and fluorescence images from left to right, three different stages during cryo-FIB milling are presented: the beginning stage (left), the middle stage with one side rough milled (middle) and the final stage with two sides fine milled (right). The fluorescence signals from two centrioles are indicated by red and orange arrows and monitored in real time. A line profile of the intensity of the fluorescence signal is correspondingly plotted below for the three stages. A schematic diagram of the thickness of the cryo-lamella at different stages is shown at the bottom. (**d**) The prepared cryo-lamella is transferred into a cryo-transmission electron microscope for cryo-ET data collection. Top, cryo-EM micrographs of the cryo-lamella at low (3600X, left) and high (33000X, right) magnifications. Bottom, a zoomed-in view of the cryo-EM micrograph at high magnification to show the centrioles, which are labelled with orange and red arrows.

First, we selected HeLa cells expressing mCherry fluorescent protein-labelled pericentrin, which is an integral component of the centrosome located in the PCM region^47^. The fluorescence signal of mCherry fluorescent protein was used to navigate cryo-FIB milling to the target centrosome. The fluorescence-labelled cells were seeded onto a custom-designed ultraviolet-sterilized OD grid and subjected to cryo-vitrification by plunge freezing. Then, the cryo-vitrified grid was assembled into an FEI AutoGrid and further loaded onto a commercial cryo-EM multiholder (Gatan, USA), ready for the subsequent cryo-transfer (Fig. 2a). Notably, there is one straight bar in the OD grid that can be used as an orientation indicator. During loading of the grid onto the cryo-holder, we adjusted the orientation of the grid to ensure that this straight bar was parallel to the holder tilt axis and therefore perpendicular to the subsequent cryo-FIB milling direction. Then, in the subsequent cryo-ET data collection, we loaded the AutoGrid and adjusted its orientation to ensure that this bar was parallel to the tilt axis.

Second and optionally, we utilized our previously developed cryo-fluorescence microscope HOPE system^28^ or SIM-HOPE system (to be submitted) to screen the cryo-vitrified cells (Fig. 2b). Both bright-field and fluorescence images under cryogenic conditions were recorded to check the thickness of the specimen and the intensity of fluorescence signals. The coordinates of the selected cells were recorded onto the map of the grid, which was used in the subsequent experiment. This step can be skipped if we can ensure that the specimen is well vitrified and contains well-distributed fluorescence signals. Notably, the cryo-transfer between our HOPE system and the subsequent ELI-TriScope system is contactless for the specimen, minimizing the risk of specimen deformation, devitrification and ice contamination.

Third, the cryo-holder with the cryo-vitrified specimen was loaded into the ELI-TriScope system. At the beginning, the cryo-holder was tilted to 30 degrees to allow the cryo-grid to face the gas injection system (GIS) (Extended Data Fig. 1c), and the predefined positions on the grid were coated with a protective organometallic layer (Extended Data Fig. 1d). Then, the holder was tilted back to -20 degrees, allowing the objective lens of the cryo-STAR system to rise to the focus position (Extended Data Fig. 1e).

Fourth, FIB and FM images were acquired with magnifications of 2,500 and 100, respectively (Fig. 2c). The target position was identified according to the fluorescence signals in the FM image with the reference coordinates recorded in the above cryo-FM screening (optional). Then, the stage was controlled to move the target position to the crossover point of the ion and electron beams, and the position of the objective lens of the cryo-STAR system could be further optimized for the best focus. Both FIB and FM images were acquired again. With good precalibration of the ELI-TriScope system (see Methods), the fluorescence signal (two adjacent bright spots for two centrioles near the cell nucleus) in the FM image can be well matched to the cell feature in the FIB image. Then, the coarse milling process was performed with a large beam current of 0.23∼0.43 nA around the target position, monitored by real-time fluorescence imaging with the cryo-STAR system.

The drift of the specimen was detected in the fluorescence image and corrected by precisely controlling the stage. When the thickness of the cryo-lamella was less than 3 *μ*m, fine milling was performed with a smaller beam current of 40∼80 pA. During the milling process, the fluorescence signal was always kept in the centre of the cryo-lamella, and the milling position was accordingly adjusted. At the beginning of milling, the two centrioles showed two strong adjacent fluorescence peaks with their intensities accurately monitored (Fig. 2c). The start of the decrease in their intensity indicated that the milling reached the target. Then, the milling process should be stopped and started again on another side of the cryo-lamella. Finally, the cryo-lamella was trimmed to less than 200 nm thickness by maintaining a sufficiently strong fluorescence intensity. Because the centriole has an overall size larger than 200 nm, the final fluorescence intensity decayed in comparison with the original fluorescence intensity.

After precise cryo-FIB fabrication, the cryo-lamella containing the target centrioles can be transferred to a cryo-electron microscope for subsequent cryo-ET data collection and *in situ* structural study (Fig. 2d).

In comparison with other reported cryo-CLEM techniques, including both nonintegrated^29,30,35^ and integrated^39,48^ workflows, ELI-TriScope simplifies the specimen transfer steps and reduces the time cost to prepare one cryo-lamella from 2∼2.5 hrs per cryo-lamella to ∼ 0.8 hrs per lamella (Extended Data Fig. 2). In addition, the precision and success rate of ELI-TriScope are also significantly improved.

### *In situ* structure of the human centriole

Human centrioles are closely related to tumorigenesis and multiple hereditary diseases^49,50^. Revealing the centriole assembly details and the roles of each centrosomal component is important. The centrioles are of variable size among species, while in mammalian cells, typical mature centrioles are approximately 230 nm in diameter and 500 nm in length^51^. Nine sets of MT triplets (MTTs) are arranged in 9-fold symmetry, and each MTT contains a full MT (A-tubule) and two partial MTs (B-tubule and C-tubule). Centrioles are polarized along their longitudinal axis, possessing different structures at the proximal and distal ends. At the proximal end, the cartwheel acts as a determinant scaffold for centriole symmetry^52^ but is degraded in mature human centrioles^53,54^. At the distal end, there are distal appendages (DAs) for mediating the attachment of ciliary vesicles to centrioles during ciliogenesis and subdistal appendages (SDAs) for positioning centrioles and cilia by anchoring MTTs^55-57^.

In this study, we spent 5 days preparing 61 cryo-lamellae of HeLa cells using our ELI-TriScope system, and all the cryo-lamellae successfully targeted human centrioles. We collected and aligned 59 cryo-ET tilt series containing 77 centrioles (Extended Data Table 1). The average thickness of our processed cryo-lamellae was 169 nm ± 43 nm, which is smaller than the narrowest part of the human centriole (Extended Data Fig. 3a). Therefore, in each tomogram, only partial centrioles exist. In some tomogram slices, we observed two new-born procentrioles next to the mature centrioles, indicating that these cells should be in the S phase of the cell cycle (S^1^) and that the centrosomes are under replicating conditions (Extended Data Fig. 3c and 3d). We selected 46 well-aligned tilt series and 55 centrioles with high integrity for further structural analysis.

Both top and side views of centrioles could be observed, and no obvious preferred orientations were found in the current dataset (Extended Data Fig. 3b), which avoids anisotropy reconstruction in the following data processing. All subvolumes of centrioles were cropped from the original tomograms along the axis from the proximal end to the distal end at 4 nm intervals and then aligned to different local regions using the subtomogram averaging (STA) approach (Extended Data Fig. 4 and Extended Data Table 1).

The human centriole shows a regular 9-fold symmetry structure in both the tomogram slice and STA reconstruction (Fig. 3 and Supplementary Video 3), which was reported to be due to the strict regulation of the highly conserved SAS-6^45,51,52,54^. Almost all the walls of centrioles are formed by MTTs, but in the very distal end of the centriole, MT doublets (MTDs) (A- and B-tubules) are observed, while the C-tubule vanishes (Extended Data Fig. 5). For the MTTs, according to the unbiased data processing and 3D classifications, approximately three-quarters of MTTs have a complete C-tubule containing 10 protofilaments and sharing 3 protofilaments (5^th^, 6^th^ and 7^th^) with the B-tubule, while a quarter of the MTTs have an incomplete C-tubule containing only approximately 5 protofilaments (Fig. 3a and Extended Data Fig. 6). Based on the relative positions of these two elements, MTTs with a complete C-tubule are located nearer the proximal end of the centriole, and those with an incomplete C-tubule are closer to the distal end (Extended Data Fig. 7), which is consistent with previous reports^58^.

**Figure 3.**
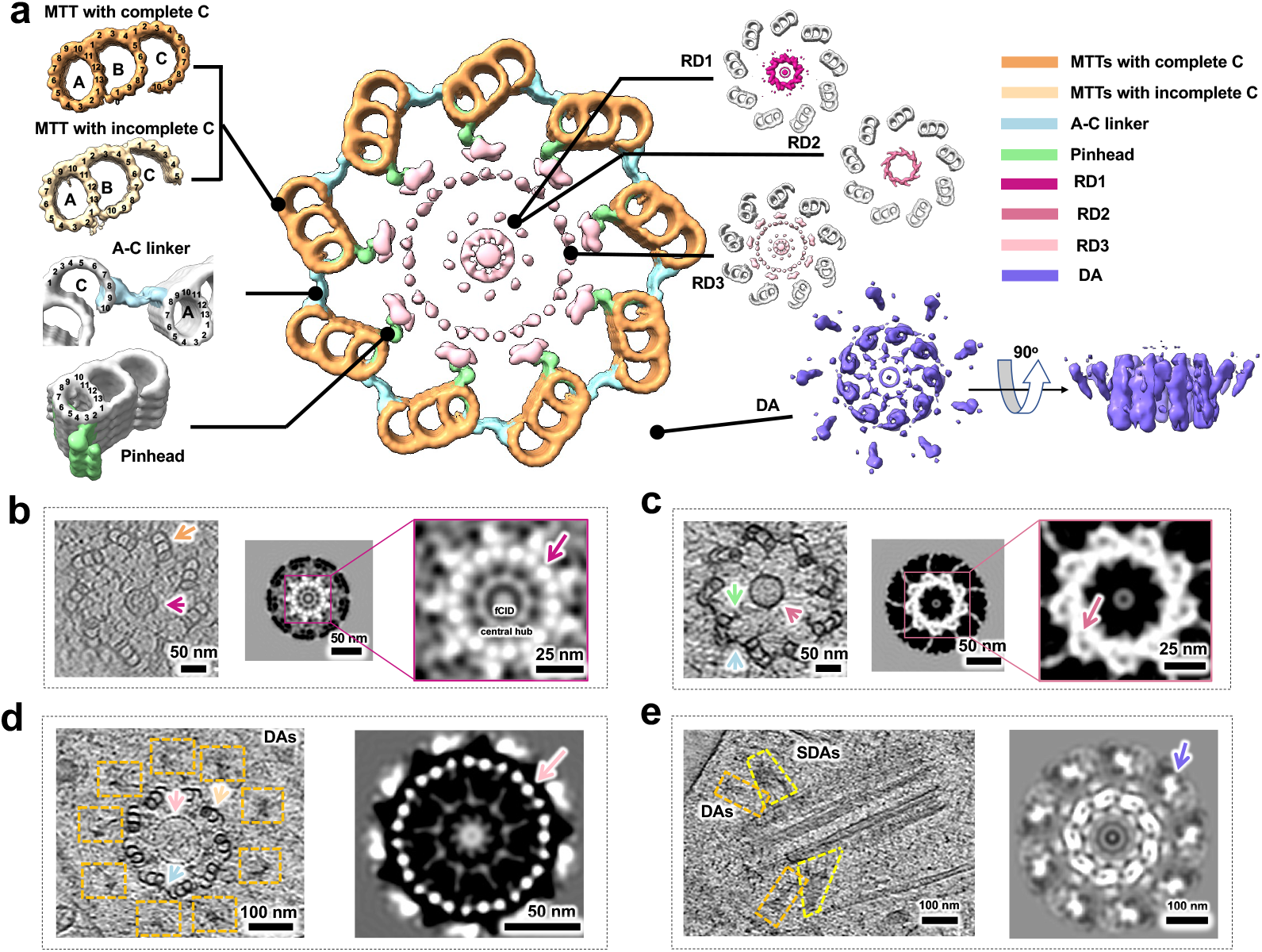
*In situ* structure of the human centriole. **(a)** Cross-sectional view of human centriole slices in different local regions, which are reconstructed by STA. The structure in each local region is correspondingly magnified and labelled. MTTs with a complete C-tubule and an incomplete C-tubule are coloured sandy brown and Navajo white, respectively. The A-C linker is coloured light blue; the pinhead is coloured light green; RD1 is coloured medium violet red; RD2 is coloured pale violet red; RD3 is coloured pink; and the DA is coloured medium slate blue. MTT, microtubule triplet; RD, ring density; and DA, distal appendage. **(b)** RD1 and MTTs with a complete C-tubule indicated by the arrows in the tomogram (left), and a cross-sectional view of the RD1 map obtained by STA (right). The RD1 peak is indicated by the arrow. The CID and CH are observed inside. **(c)** RD2, A-C linker and pinhead indicated by the arrows in the tomogram (left), and cross-sectional view of the RD2 map obtained by STA (right). The RD2 peak is indicated by the arrow. **(d)** RD3, MTTs with an incomplete C-tubule, A-C linker and DA indicated by arrows and boxes in the tomogram (left), and cross-sectional view of the RD3 map obtained by STA (right). The 27 rod-like density peaks of RD3 are indicated by the arrow. **(e)** SDA and DA indicated by boxes in the tomogram (left), and cross-sectional view of the DA map obtained by STA (right). The baseball-bat-shaped density of the DA is indicated by the arrow. The colour schemes in (**b-e)** are the same as that in (**a)**.

The diameter of the centriole distal end where MTDs emerge is ∼220 nm (Extended Data Fig. 8a), while the regions in the same centriole where MTTs with an incomplete C-tubule exist have an average diameter of ∼240 nm (Extended Data Fig. 8b). When MTTs with a complete C-tubule exist, centrioles show a larger diameter of ∼270 nm (Extended Data Fig. 8c). Our observation that the human centriole has the narrowest diameter in the distal region is consistent with previous studies^51,59^.

According to the STA reconstructions of MTTs and single A-tubules (∼25 Å resolution), the MT walls of the human centriole have an ∼8.4 nm repeating periodicity (Extended Data Fig. 6), similar to that in *Paramecium tetraurelia* (*P. tetraurelia*), *Chlamydomonas reinhardtii* (*C. reinhardtii*), and centrioles isolated from *Homo sapiens* (*H. sapiens*) KE37 cells^60^. For MTTs with a complete C-tubule, a linker density can be observed between the A- and C-tubules from adjacent MTTs, referred to as the A-C linker (Fig. 3). In the STA reconstructions of the A-C linker, we found that it binds to the 8^th^ protofilament of the A-tubule in one MTT and the 8^th^ and 9^th^ protofilaments of the C-tubule in the adjacent MTT (Fig. 3a), consistent with previous reports on *Trichonympha agilis* (*T. agilis*) and Chinese hamster ovary (CHO) cells^51,61^. This shows that the A-C linker conformation is highly conserved among species. For those MTTs with an incomplete C-tubule, the A-C linker density is missing in some regions but remains between the A-tubule and the edge of the incomplete C-tubule (Fig. 3c and Fig. 3d). It seems that even for an incomplete C-tubule with only approximately 5 protofilaments, the A-C linker can still connect it to the A-tubule of the adjacent MTT. Possibly, the A-C linker connects with other protofilaments in the incomplete C-tubule.

In addition to the A-C linker, another important component that interacts with the MTT is the pinhead. The pinhead has been reported to extend from the MT wall to bridge the A-tubule and the central cartwheel in the proximal end of the centriole^62,63^. However, whether the pinhead exists in the mature centriole of humans remains unknown. In this study, we observed the pinhead density in most centrioles (Fig. 3c). It contains at least two fibrous components and interacts with the 3^rd^ protofilament of the A-tubule (Fig. 3a). The remaining part of the pinhead that links the central cartwheel is highly dynamic; it can only be traced in some tomogram slices but is averaged out in the STA reconstruction.

### Internal and external structures of human centrioles

Precise assembly of the centriole and maintenance of the cohesion stability require internal scaffold structures. The c<u>artwheel</u> was the first observed internal structure and believed to be the scaffold for centriole biogenesis as well as the determinant for the centriole symmetry, which has been studied in several species, such as *P. tetraurelia* and *C. reinharadtii*^52,54,62,64^. In the human centriole, we observed several types of ring structures with different shapes and diameters inside MTTs. After 3D classification with 9-fold symmetry applied, three major populations were identified, named ring density (RD)1, RD2 and RD3, with diameters of approximately 50 nm, 65 nm and 100 nm, respectively (Fig. 3a).

Cartwheel inner density (CID) (∼ 7 nm in diameter), central hub (CH) (∼ 25 nm in diameter) and spoke structures were reported to exist in the middle of centrioles in the proximal region^54^. In this study, we found similar structures in RD1 from the STA map and some raw tomograms (Fig. 3a and b) but not in all the RD1-existing centrioles. RD1 is probably a cartwheel-related structure, but its internal components may change during the development stage of centrioles. The 9 density peaks of RD1 suggest that it may form a rod-like structure. Unlike the reported longitudinal periodicity of ∼4.2 nm along the CH and ∼8.1 nm along the CID from isolated centrioles of *H. sapiens*^54^, we did not find similar periodicity of RD1 in our STA analysis.

The diameter of RD2 is slightly larger than that of RD1, and no structural densities were found within RD2 (Fig. 3c). In the subtomogram averaged volume with 9-fold symmetry, we observed a blurred density of the RD2 wall in comparison with the peak density of the RD1 wall (Fig. 3c), suggesting that RD2 might have different symmetry than 9-fold or have high dynamics.

RD3 is another internal structure of the human centriole observed in our tomograms, and to the best of our knowledge, it has not been discussed before. It has a diameter of 100 nm and contains 27 peak densities along the wall (Fig. 3d and Supplementary Video 3). It has no obvious inner structure and is located near the pinhead region. Interestingly, this RD3 structure is often associated with MTTs possessing an incomplete C-tubule and a DA (Fig. 3d and Supplementary Video 3), indicating that RD3 is located at the distal end of the centriole and may be related to DA components. In the STA volume with 9-fold symmetry, RD3 shows 27 clear rod-like structures, suggesting that it may play a role in maintaining the cohesion of MTTs in the distal end region. Meanwhile, we noted an RD3-like density in a previous study of centrioles using the plastic section of HeLa cells^55^; however, it was not carefully and explicitly studied.

DA is a typical structure of mature centrioles in many species, and it is also observed in our tomogram slice of the human centriole (Fig. 3e and Supplementary Video 4). Each DA protrusion connects to one MTT and extends towards the outward and distal ends of centrioles, whose terminus has a diameter of ∼450 nm. In the subtomogram averaged volume with 9-fold symmetry, the DA shows a baseball-bat-shaped structure of each protrusion (Fig. 3a). Interestingly, in this STA volume, we also observed a clear 100 nm diameter RD3 density that connects to the pinhead of MTTs inside the centriole. For the SDA, it can be found near the DA in the tomogram slice (Fig. 3e and Supplementary Video 4) but is averaged out in the STA reconstruction.

## Discussion

In this study, to achieve precise cryo-FIB fabrication of cells and efficiently prepare target cryo-lamellae for subsequent cryo-ET *in situ* structural study, we developed an ELI-TriScope system by incorporating a cryogenic fluorescence simultaneous monitor system (cryo-STAR) into a commercial dual-beam SEM. To enable the electron beam, ion beam and light beam to be focused on the same coincidence point, a new cryo-transfer system utilizing a commercial cryo-holder was designed to replace the conventional chamber door of the dual-beam SEM, which allows the objective lens of cryo-STAR to be positioned just underneath the cryo-specimen. Our cryo-transfer system achieves seamless and contactless cryo-specimen transfer among our HOPE cryo-FM system, ELI-TriScope system and FEI Titan Krios cryo-electron microscope and largely decreases the risk of specimen damage, deformation, devitrification, and ice contamination.

In the ELI-TriScope system, after careful precalibration, the electron beam, ion beam and light beam are focused on the same point, and then, the cryo-specimen can be imaged simultaneously by the FIB beam and light beam with the same field of view. Therefore, cryo-FIB milling can be precisely navigated by monitoring the real-time fluorescence signal, resulting in accurate cryo-fabrication of cells in the target region. Compared to previously reported cryo-CLEM workflows for cryo-lamella preparation, our ELI-TriScope technique does not require prespreading of fluorescence beads in the sample, avoids the sophisticated correlation procedure between cryo-FM and cryo-FIB images, and simplifies the overall workflow of site-specific cryo-lamella preparation with a high success rate.

In the present work, we applied our ELI-TriScope technique to study the *in situ* structure of the human centriole in HeLa cells, which is a challenging task using previous cryo-CLEM techniques. Recent advances in cryo-EM have greatly promoted the structural study of centrioles in *C. reinhardtii, P. tetraurelia, Naegleria gruberi* (*N. gruberi*), *H. sapiens*, etc.^51,54,58,60^. However, great difficulties are still encountered in the study of high-resolution structures of centrioles in their native environment. For *C. reinhardtii*, the exact location of the centriole can be tracked along the flagella, but the location of centrioles in most other species cannot be easily targeted, leading to a significantly low efficiency of cryo-FIB milling. This greatly limits the *in situ* structural study of centrioles in mammalian cells.

Using ELI-TriScope, we were pleased to observe that we could prepare 61 high-quality cryo-lamellae of HeLa cells within 5 days and that all the cryo-lamellae could target the centrioles accurately and successfully. We were therefore able to collect a large amount of cryo-ET tilt series of cryo-lamellae and study the *in situ* structure of the human centriole. We observed multiple components of human centrioles in their native states. With a preliminary STA procedure, we could resolve the single protofilaments of MTTs and observe their longitudinal periodicity.

With the 9-fold geometry in the centriole of almost all organisms, the MTs that comprise the centriole cylinder can be singlet, doublet or triplet depending on species. It has been illustrated that centrioles in early *Caenorhabditis elegans* (*C. elegans*) embryos have MT singlet (MTS) assembly^65^. In addition, centrioles from *Drosophila melanogaster* S2 cells are composed of a mixture of singlets and doublets^51^, whereas those in sperm cells are uniquely long and consist of MTTs^66^. In addition, previous studies showed that some species, including *C. reinhardtii* and human KE37 cells, have MTTs in the proximal region and MTDs in the distal ends^58,60^. Here, we found that human centrioles in HeLa cells are mostly composed of MTTs and only have MTDs at the very distal end region. Moreover, we also observed many internal and external densities of human centrioles, including RD1, RD2, RD3 and DAs. Besides aligning with previous reports, the existence of RD1∼3 in the human centriole has not been reported before. We found that RD1 is associated with the cartwheel in the proximal region, and RD3 is associated with the DA and is located around the distal region. In addition, a helical inner scaffold structure was found to maintain MTT cohesion under compressive force^60^, and it can also be traced in the distal regions of centrioles in our dataset (Supplementary Video 3). These discoveries of ours will be further studied by collecting more cryo-ET data and improving the completeness and resolution of STA in the future.

Overall, we developed an advanced precise and efficient cryo-FIB fabrication technique with the name ELI-TriScope and applied this technique to study the 3D *in situ* structure of the human centriole for the first time to our knowledge. Our results have demonstrated that ELI-TriScope will have wide application in future *in situ* structural biology and in studying the high-resolution ultrastructure of specific events in the cell.

## Methods

### Incorporating the cryo-STAR system into a dual-beam SEM

To build the cryo-STAR system (Fig. 1a), an embedded widefield optical imaging system was mounted onto an existing port of a commercial dual-beam SEM (FEI Helios 600i, ThermoFisher Scientific, USA). The light path occupies a large port, and the control cables use a small port. In the vacuum chamber, a high-NA dry objective lens (LMRlanFL N 100X, NA/WD: 0.8/3 mm, Olympus, Japan) was placed underneath the vitrified specimen, opposite to the pole pieces of the electron and ion columns. It was installed with an angle of 18 degrees to the electron beam, nearly perpendicular to the grid plane. The position of the objective lens can be finely adjusted in three dimensions using an XYZ piezo stage (Micronix, USA) during focusing.

Cryo-STAR is equipped with an epifluorescence system with a full-spectrum (DAPI/GFP/RFP/Cy5) white LED light source (FC904s, Shanghai Fluoca Technology Co., Ltd., China). The beam excited from the white LED light source is expanded to a diameter of 25 mm, is reflected by a dichroic mirror, and passes through into the vacuum chamber. Then, the excitation beam is reflected to the objective lens by a reflector. The emission light from the sample goes back along the incident illumination path and passes through the dichroic mirror and the emission filter. The final image is detected by a high-sensitivity CMOS camera (Moment, Teledyne Photometrics, USA). Micro-Manager (ver. 2.0)^67^ was used to control the camera and record fluorescence images. All lenses used in the system were chosen as achromatic doublets. Optical apertures and filters were used to improve the illumination quality.

### Calibration and operation of the cryo-STAR system

The cryo-STAR system requires precalibration by a finder or index grid with obvious features, such as fluorescent beads. For example, a finder grid is transferred into the SEM chamber by a holder, and then, an obvious feature on the grid is found and adjusted to the crossover point of the electron and ion beams. After centring this feature in both the electron and ion images, the position of the objective lens is adjusted to make the feature clear and at the centre of the image. In this way, the light beam, electron beam and ion beam are adjusted to be at the same focal point, normally with a position accuracy better than 1 μm. The calibration parameters can be checked again during the cryo-FIB milling process to ensure the precision.

In the monitoring process, to avoid potential warming up of the cryo-vitrified specimen and induction of devitrification, a stroboscopic exposure mode was used with each exposure time of 200 ms and a camera frame rate of 1-0.5 fps. The total illumination power was controlled at ∼ 20 μW with a minimum illumination intensity of 0.05 W/cm^2^. Compared to other cryo-FM techniques, such as cryo-iPALM^37^, which always require acquisition of raw image sets with 25,000 to 75,000 frames or more at a minimum laser power of ∼ 1 W and a minimum illumination intensity of 0.1-1 kW/cm^2^, the heating effect induced by our cryo-STAR illumination is minimal, which largely avoids the risk of devitrification of cryo-specimens. Indeed, we never observed the phenomena of warming up of the frozen specimen and bleaching of the fluorescence signal in our studies.

### Cryo-transfer into ELI-TriScope

Before sample loading, the multispecimen cryo-holder (Gatan, USA) with a homemade tip was cooled to liquid nitrogen (LN_2_) temperature. Then, the AutoGrid with the sample was mounted into the tip in the cryo-holder workstation. The position of the objective lens of cryo-STAR was reset to ensure that there was sufficient space above for sample loading. Then, the cryo-holder was inserted into the vacuum transfer system with prepump for approximately 60 seconds. Then, the mini-gate valve was manually opened to allow the cryo-holder to be inserted into ELI-TriScope. After waiting 5 minutes to recover the high vacuum of the chamber and confirming that the tip of the cryo-holder was far from the pole pieces of the electron and ion columns, the cryo-holder was tilted to 30 degrees right in front of the GIS^48^. The predefined sample position on the grid was coated with a protective organometallic layer for ∼ 5 s. Then, the cryo-holder was tilted back to -20 degrees, allowing the objective lens of the cryo-STAR system to rise to the focus position for subsequent real-time fluorescence-navigated cryo-FIB milling.

### Culture of HeLa cells and vitrification

HeLa cells expressing mCherry fluorescent protein-labelled pericentrin^47^ were seeded onto ultraviolet-sterilized OD grids (T11012SS, TIANLD, China) and cultured in complete DMEM supplemented with 10% foetal bovine serum, 1% penicillin and streptomycin. After 48 hrs of culture with 5% CO_2_ at 37 °C, the grids were subjected to plunge freezing by backside blotting and vitrification using a Leica EM GP1 (Leica Microsystems, Germany). Then, the cryo-vitrified grid was assembled into an FEI AutoGrid (ThermoFisher Scientific, USA) and further loaded onto a commercial cryo-EM multiholder (Gatan, USA), ready for the subsequent cryo-transfer.

### Cryo-ET data collection

All cryo-ET data were acquired on an FEI Titan Krios cryo-TEM (ThermoFisher Scientific, USA) equipped with an energy filter and a K2 Summit Direct electron detector (DED) (Gatan, USA) using the SerialEM package^68^ at 300 kV. Microscope operation and filter tuning adjustment were performed using SerialEM and Digital Micrograph (Gatan, USA). The slit width of the filter was set to 40 eV. Tilt series were acquired with a unidirectional tilt scheme ranging from -45 to 45 degrees with a step of 2 degrees, and the target defocus was set to -6 μm. The nominal magnification was 33,000 ×, resulting in a pixel size of 4.3 Å. Individual tilt images were acquired as 3838 × 3710 pixels movies including 10-12 frames. All procedures for cryo-EM imaging were performed under low-dose conditions.

### Data processing

Image movies were processed by motion correction and CTF estimation in Warp^69^. The produced tilt series were aligned using IMOD^70^ and then transferred back into Warp to reconstruct full tomograms. The filament model in Dynamo^71^ was used to manually pick 55 centrioles in 46 tilt series from the proximal end to the distal end, which was judged by the clockwise or counterclockwise direction of MTTs in the cross-sectional view of the centriole^72^. Every particle in the same centriole was separated by 4 nm. All the subvolumes were cropped in Warp, and the subsequent data processing, including 3D classification, autorefinement, and duplicate removal, was performed in Relion^73^. Different components of the centriole were aligned and refined by shifting the volume centre to different local regions (Extended Data Fig. 4). To achieve unbiased reconstructions, the templates used in the alignment procedures were all data-driven references, and none of the other reported structures were used in this study.

### Segmentation and visualization

Imaris 9.8.0 (Oxford Instruments, UK) was applied to segment centrioles from 2 binned tomograms using a SIRT-like filter equivalent to 15 iterations. The typical characteristics of the centriole were labelled manually and coloured differently. All figures were created using ChimeraX^74^, and all movies were generated in Imaris 9.8.0.

## EXTENDED DATA

**Extended Data Fig. 1.**
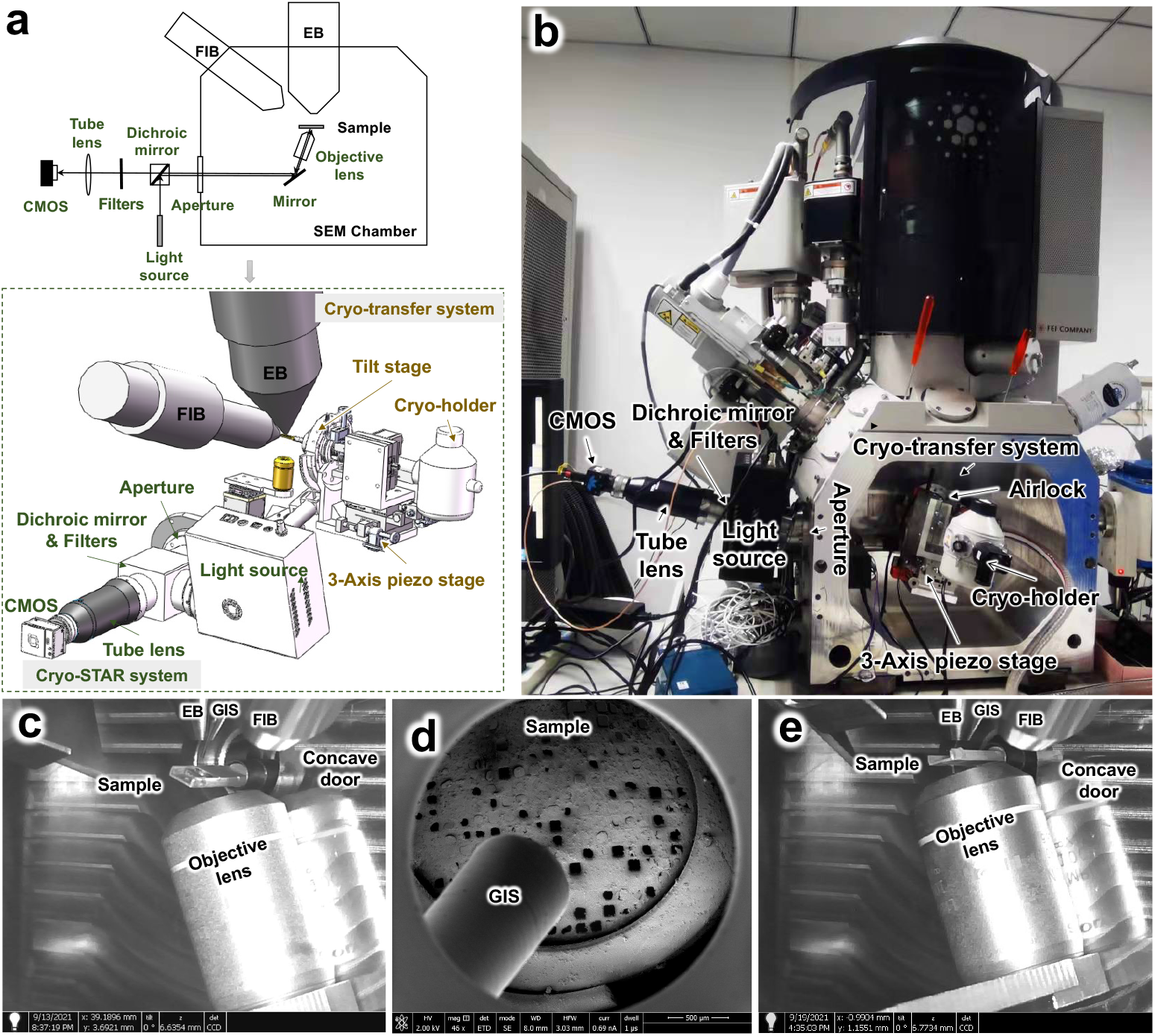
Design and construction of ELI-TriScope. (**a**) Schematic diagram of the ELI-TriScope system showing two major components, a cryo-holder-based cryo-transfer system and a cryo-STAR system. Along the direction of the optical path of the cryo-STAR system, the excitation light beam coming out of the light source is reflected by a dichroic mirror into the aperture, reflected into the objective lens by an adjustable mirror, and then excites the fluorescent molecules in the cryo-specimen. Next, the excited fluorescence is received by the objective lens, is reflected into the aperture by the adjustable mirror, passes through a dichroic mirror, is filtered and is focused on a CMOS camera through a tube lens. The cryo-transfer system is equipped with a 3D motorized device that is adapted to the cryo-holder. (**b**) Photograph of our built ELI-TriScope system based on FEI Helios NanoLab 600i. A custom-designed cryo-transfer system replaces the original chamber door, and the cryo-STAR system is fixed on the other side of the chamber. (**c**) Photograph of the internal architecture of ELI-TriScope. The cryo-holder is tilted 30 degrees to the right to face the GIS for coating. (**d**) Observation of the coating status by SEM imaging at 40X magnification after GIS coating. (**e**) Position for cryo-FIB fabrication. The cryo-holder is tilted to -15 degrees. The objective lens of cryo-STAR rises to the focus position. Each part of the system is indicated and labelled. EB, electron beam. GIS, gas injection system.

**Extended Data Fig. 2.**
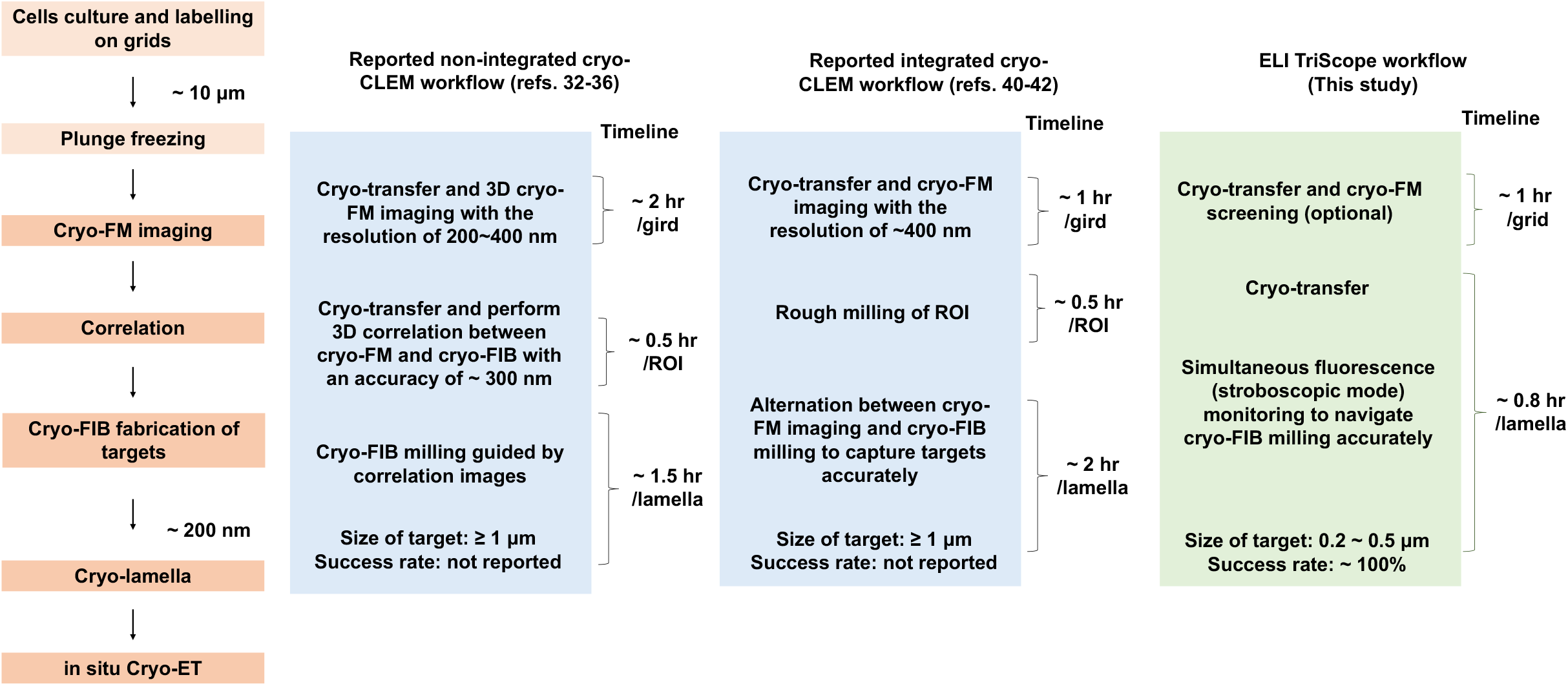
Comparison of three different cryo-CLEM workflows for site-specific cryo-FIB fabrication. A conventional experimental procedure of the *in situ* cryo-ET study is described (left), including steps from cell culture to plunge freezing, cryo-FM imaging, cryo-FIB imaging and correlation, cryo-FIB milling, production of a cryo-lamella and cryo-ET data collection. The reported cryo-CLEM workflows from cryo-FM imaging to cryo-FIB milling can be classified into nonintegrated workflows^29,30,35^ and integrated workflows^39,48^. In a nonintegrated workflow, the cryo-FM and cryo-FIB steps are carried out individually and sequentially. Then, with the help of fiducial markers, cryo-FM images and cryo-FIB images are aligned by 3D correlation software and used to guide cryo-FIB milling. In an integrated workflow, such as in the commercial products iFLM (FEI) and METEOR (Delmic), due to the limited space inside the chamber and the limitation set by the cryo-stage, only a low-NA objective lens can be installed just alongside the ion column. Cryo-FM images are alternatively acquired before and after cryo-FIB milling to check the target fluorescence signal and accurately guide cryo-FIB milling, which reduces the overall throughput of cryo-lamella production. For both nonintegrated and integrated workflows, with the cryo-FM resolution and correlation accuracy limitations, the sizes of the studied targets (e.g., mitochondria, lipid droplets) are normally larger than 1 μm, and the success rate of cryo-lamella production was not reported. In the ELI-TriScope workflow, after the optional cryo-FM imaging, there is no need for a 3D correlation step, and the fluorescence signal can be acquired in real-time stroboscopic mode to simultaneously monitor the cryo-FIB milling procedure. With this, the time cost to prepare one cryo-lamella using the ELI-TriScope workflow can be greatly reduced to ∼ 0.8 hrs per lamella, in comparison with 2∼2.5 hrs per cryo-lamella for previously reported nonintegrated and integrated workflows.

**Extended Data Fig. 3.**
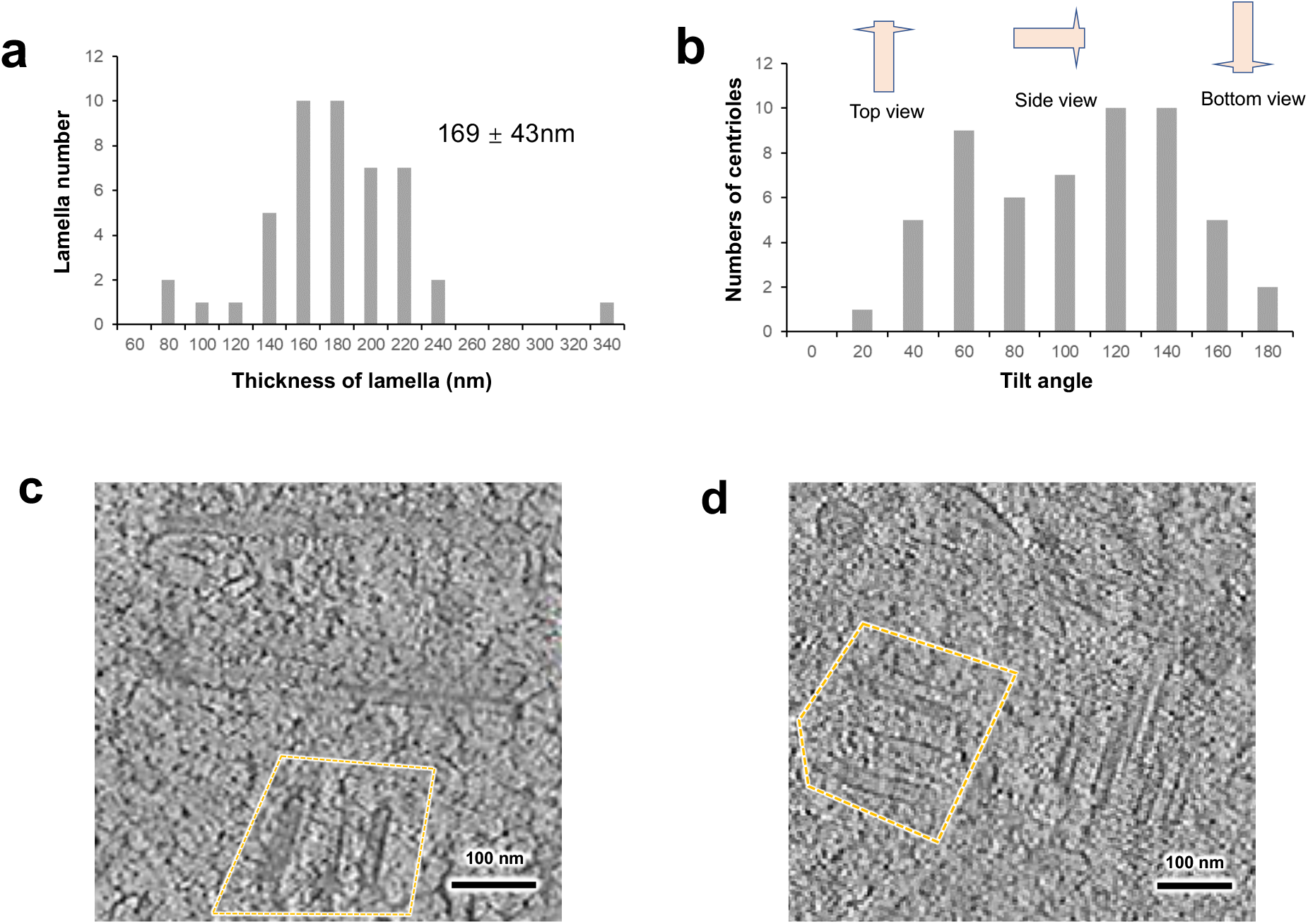
Statistical histograms of collected cryo-ET data and some typical centrioles in tomogram slices. **(a)** Statistical histogram of the lamella thickness. **(b)** Tilt angle distributions of the centrioles. **(c-d)** New-born human procentrioles. Slice views of possible new-born procentrioles (sandy brown arrows) were observed in different tomograms.

**Extended Data Fig. 4.**
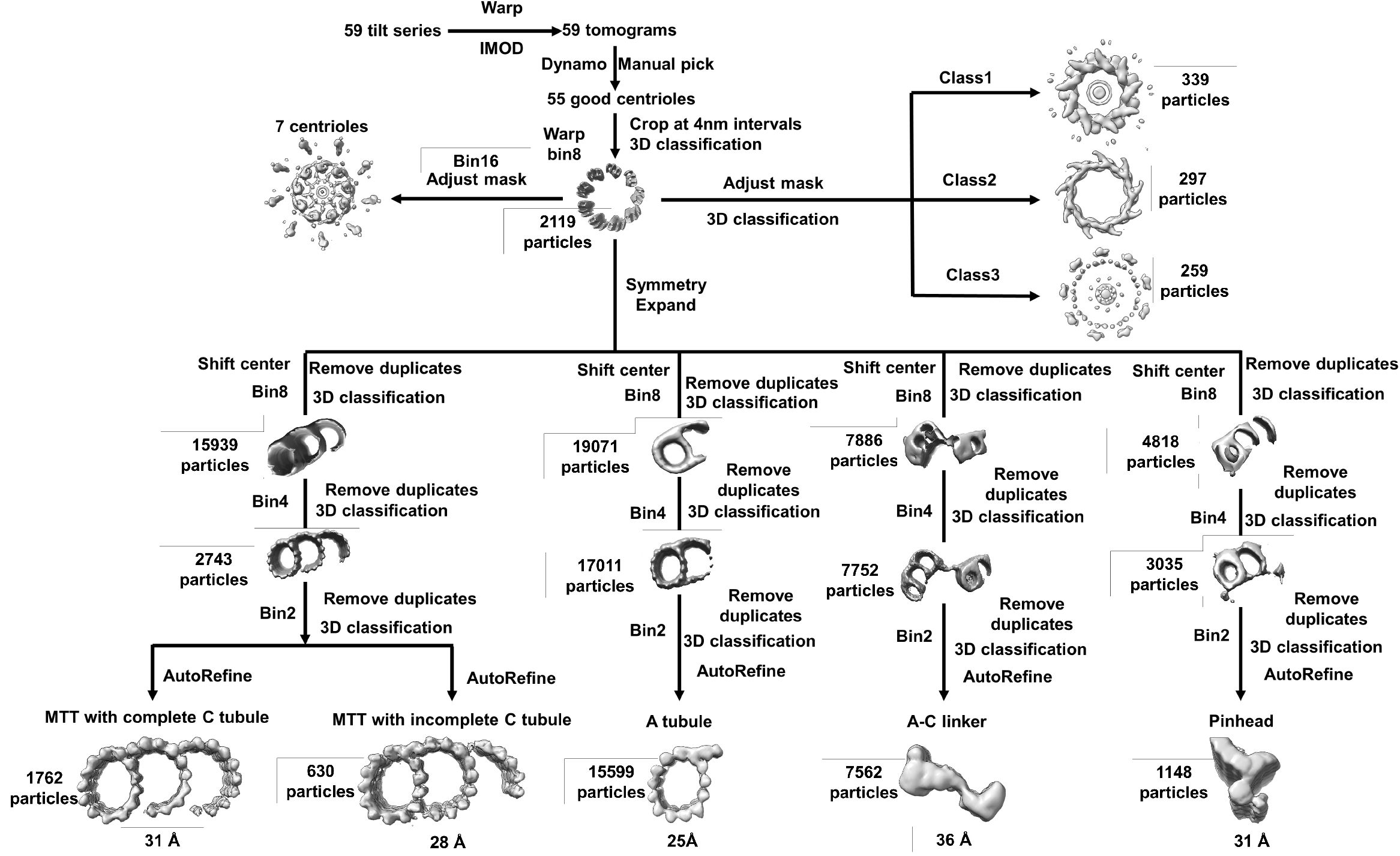
Cryo-ET data processing workflow for different parts of the human centriole.

**Extended Data Fig. 5.**
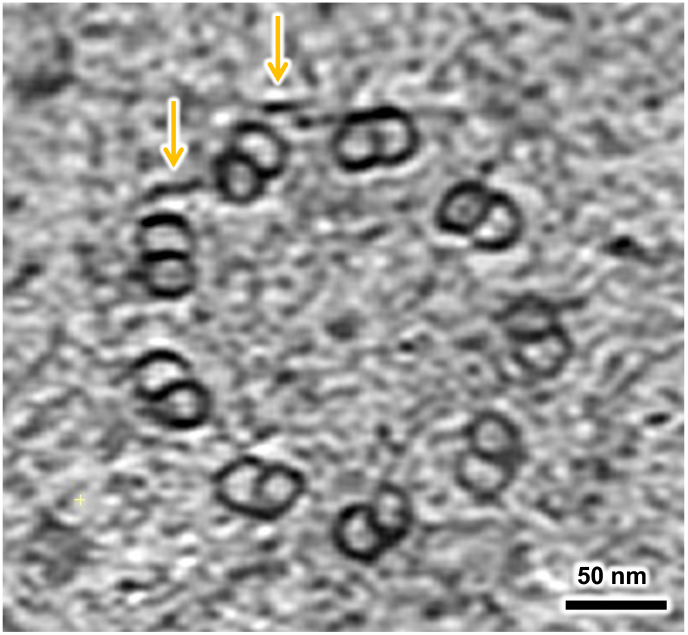
MTD structure of the human centriole in a slice view of a tomogram. The missing or incomplete C-tubules are indicated by yellow arrows.

**Extended Data Fig. 6.**
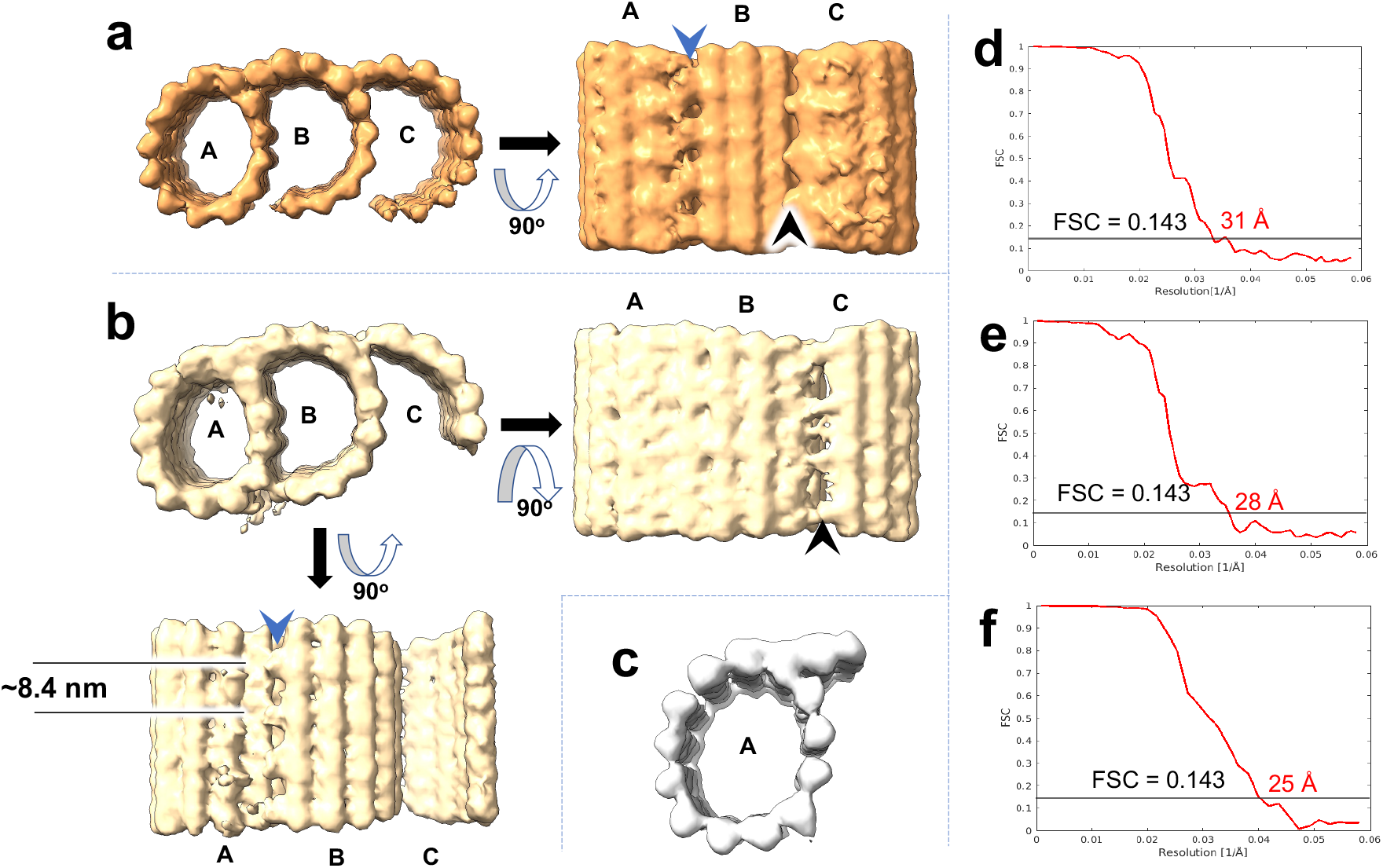
STA of MTTs from the human centriole. **(a)** Top view and side view of MTTs with a complete C-tubule, showing junctions between the A- and B-tubules (blue arrow) and B- and C-tubules (black arrow). **(b)** Top view and two side views of MTTs with an incomplete C-tubule, showing junctions between the A- and B-tubules (blue arrow) and B- and C-tubules (black arrow). **(c)** Top view of a single A-tubule. The gold-standard Fourier shell correlation (FSC) curves for the MTTs in (a), (b) and (c) are shown in (d), (e) and (f), respectively. The resolutions are calculated according to the FSC 0.143 criterion and labelled.

**Extended Data Fig. 7.**
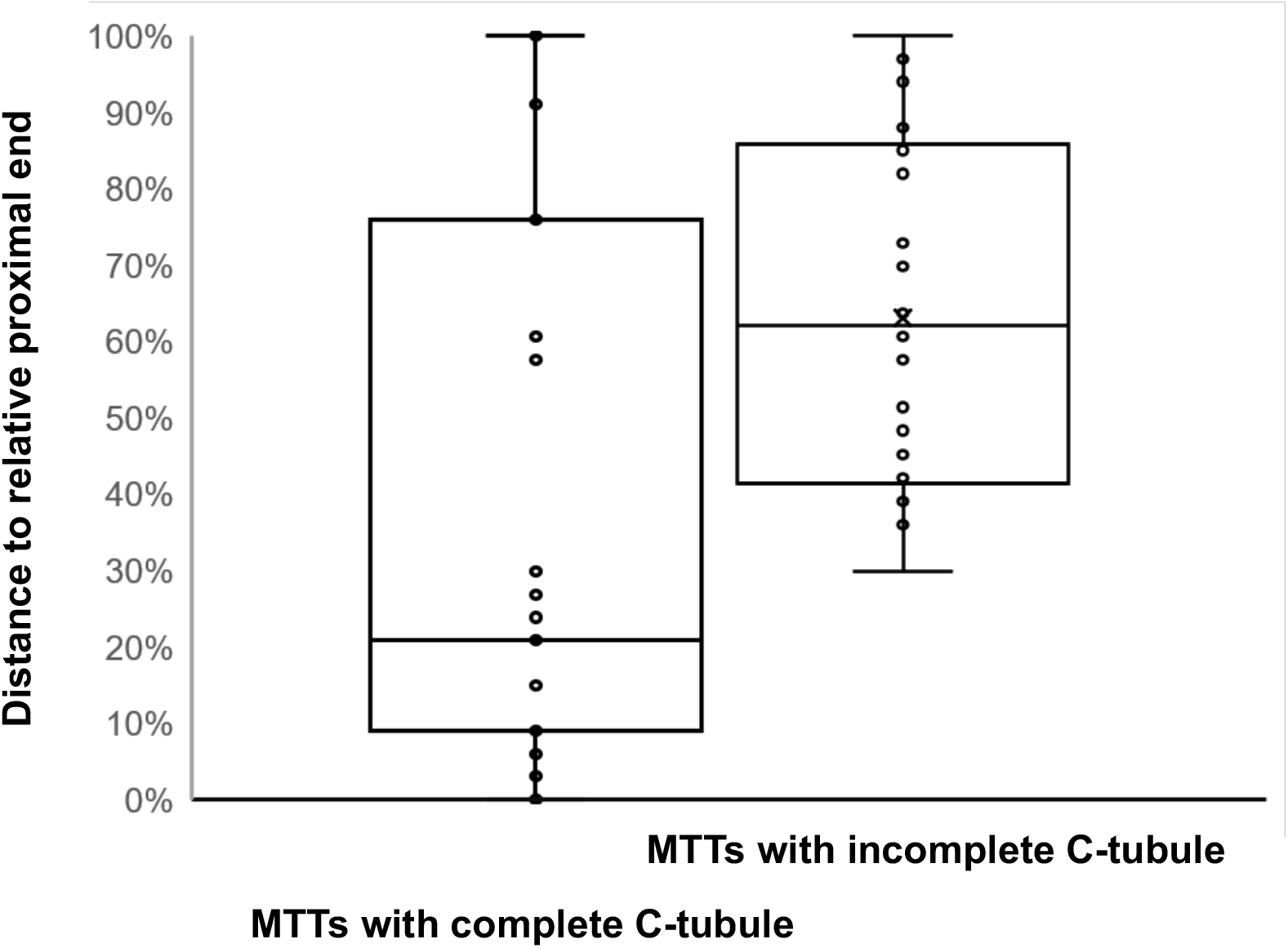
Distributions of MTTs with a complete C-tubule and an incomplete C-tubule in a single centriole. The two ends of the centriole in the tomogram are taken for the relative proximal and distal ends. The coordinates of the refined MTT elements are projected onto the central symmetry axis of the centriole to calculate the relative distances.

**Extended Data Fig. 8.**
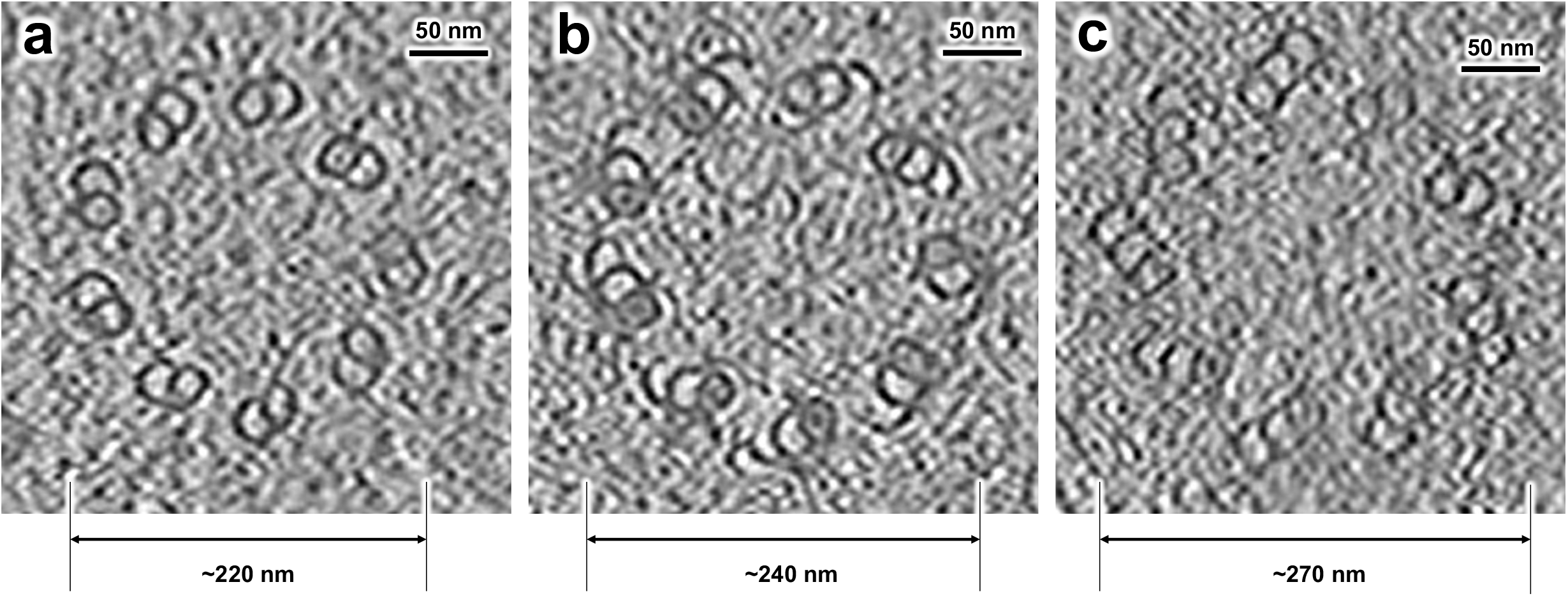
Different diameters of human centrioles in the slice view of tomograms.

**Extended Data Table 1.** Statistics of cryo-ET data collection and image processing.

| Data acquisition |  |  |  |  |  |
| --- | --- | --- | --- | --- | --- |
| Microscope | Titan Krios G2 |  |  |  |  |
| Voltage (kV) | 300 |  |  |  |  |
| Detector | Gatan K2 |  |  |  |  |
| Energy filter | Gatan GIF Quantum, 40 eV |  |  |  |  |
| Mode | Counting |  |  |  |  |
| Pixel size (Å) | 4.3 |  |  |  |  |
| Stage tilting angle | -45° - 45° |  |  |  |  |
| Number of images | 45 - 60 |  |  |  |  |
| Exposure per image(e/Å²) | 3 |  |  |  |  |
| Exposure per tilt(e/Å²) | 120 - 140 |  |  |  |  |
| Defocus range (µm) | -6 - -9 |  |  |  |  |
| Software | SerialEM |  |  |  |  |
| Sub tomogram analysis |  |  |  |  |  |
| Software | RELION-3.1/Warp |  |  |  |  |
| Data set | MTTs with<br>complete C | MTTs with<br>incomplete<br>C | A-tubule | A-C linker | pinhead |
| Number of<br>particles | 1762 | 630 | 15599 | 7562 | 1148 |
| Symmetry | C1 | C1 | C1 | C1 | C1 |
| Resolution<br>(Å) | 31 | 28 | 25 | 35.9 | 31 |
| Map pixel<br>size (Å) | 8.6 | 8.6 | 8.6 | 8.6 | 8.6 |

### Data availability

The raw tilt series used in this study has been deposited in EMPIAR (the Electron Microscopy Public Image Archive) China (http://www.emdb-china.org.cn) under accession code EMPIARC-200003. The sub-tomogram averaged cryo-EM maps of the MTTs with complete C tubule, MTTs with incomplete C tubule, A tubule, A-C linker and pinhead have been deposited in the Electron Microscopy Database (EMDB) with the accession codes EMD-33417, EMD-33418, EMD-33419, EMD-33420 and EMD-33421, respectively. Other data that support the findings of this study are available from the corresponding author upon request.

### Code availability

The LabVIEW program for device controlling is hardware-dependent and the code to control the stage of our ELI-TriScope is available at the GitHub: https://github.com/hilbertsun/ELI-TriScope.

## Acknowledgements

We would like to give our special thanks to Dr. Wei Xu for his role of mentoring the team in the field of cryo-electron microscopy. We would like to thank Dr. Fulin Wang and Prof. Jianguo Chen from Peking University for their kind help of providing Hela cells expressed mCherry fluorescent protein-pericentrin. We are grateful to Drs. Xiaojun Huang, Boling Zhu, Xujing Li, Ms. Lei Sun, Lu Qin and Mr. Yongsheng Chen for their help on cryo-ET data collection, Mr. Chen Qi, Ms. Yan Teng, Dr. Yun Feng, Ms. Qing Bian and Ms. Chunliu Liu for their help on visualization and segmentation, Dr. Ping Shan, Dr. Yanxia Jia and Ms. Shuoyuan Li for their assistance of project management. All the cryo-FM, cryo-FIB and cryo-ET work were performed in Center for Biological Imaging (CBI, http://cbi.ibp.ac.cn), Institute of Biophysics (IBP), Chinese Academy of Science (CAS).

This work was equally supported by grants from National Natural Science Foundation of China (31830020 to FS), Ministry of Science and Technology of China (2017YFA0504700 to GJ), and Chinese Academy of Sciences (XDB37040102 to FS). This work was also supported by Technological Innovation Program of Chinese Academy of Sciences (29Y8CZ021001 to GJ and 29Y7CZ041001 to JZ) and CAS Key Technical Support Personnel Project (29Y9CQ041 to GJ), and by grants from National Natural Science Foundation of China (31801199 to SL and 31801201 to XJ).

## Author contributions

FS, GJ and YZ initiated and supervised the project. GJ and SL designed and built the cryo-STAR system. GJ, JZ and SL designed and built the cryo-holder based vacuum transfer system for SEM. GJ wrote the control software. XJ and XZ performed cell culturing and vitrification. SL and GJ performed cryo-fabrications using ELI-TriScope system. SL, XJ and XZ collected cryo-ET tilt series. YZ, ZW, TN and GY performed image processing. SL, ZW, YZ, GJ and FS analyzed the data and wrote the manuscript.

## Additional information

Supplementary information

**Supplementary Video 1. Design and working principle of the ELI-TriScope system**. The video shows a 3D view of the ELI-TriScope system and the design of each component. The working principle of ELI-TriScope is shown via an animation.

**Supplementary Video 2. Whole cryo-CLEM workflow based on the ELI-TriScope system**. A detailed cryo-CLEM workflow using the ELI-TriScope system is shown from precalibration of the cryo-STAR system to cell culturing, plunge freezing, cryo-FM screening (optional), cryo-transfer, cryo-FIB milling with real-time cryo-fluorescence monitoring, and the final cryo-ET data acquisition for the target cryo-lamella.

**Supplementary Video 3. Top view of one incomplete human centriole from one tomogram**. The centriole was manually segmented from the tomogram using Imaris and highlighted. MTTs are shown in pink, RD3 in green, the inner scaffold in light blue and DAs in yellow.

**Supplementary Video 4. Side view of one incomplete human centriole from one tomogram**. The centriole was manually segmented from the tomogram using Imaris and highlighted. MTTs are shown in yellow, DAs in green, and SDAs in pink.

